# Cell-Cycle Regulation In Planta: Linking Nucleus Geometry and CYCB1;1 Marker Accumulation

**DOI:** 10.64898/2026.11.19.767918

**Authors:** T. Pasternak, I. Lebedev

## Abstract

Cell division is a fundamental biological process underlying plant growth and development and ensuring the transmission of genetic and epigenetic information. Although cell-cycle regulation has been extensively investigated using suspension-cultured cells and molecular reporters, these experimental systems do not fully reproduce the hormonal environment, cellular identity, and chromatin organization of cells within intact plant tissues. From a physical perspective, cell-cycle progression also involves substantial structural remodeling of genetic material, particularly during the transition from the G2 nuclear configuration to condensed mitotic chromosomes. The magnitude of this remodeling may depend on the initial geometry and organization of chromatin.

Here, we quantitatively characterized nuclear and chromatin architecture in different cell types of the *Arabidopsis thaliana* root, focusing on the structural differences between G2 and mitotic states. In parallel, we assessed CYCB1 reporter expression using GUS and GFP signal intensities. We found that epidermal and cortical cells exhibited greater G2-to-M structural distances accompanied by higher CYCB1 reporter signals, whereas inner root tissues showed smaller structural distances and lower reporter intensities.

These findings reveal a relationship between cell-type-specific chromatin architecture and CYCB1 reporter accumulation. We propose that differences in the structural remodeling required for mitotic entry contribute to the spatial heterogeneity of cell-cycle regulation in plant tissues. Importantly, CYCB1 reporter intensity should not be interpreted as a direct or universal quantitative measure of cell-cycle activity across structurally distinct cell types. Instead, quantitative analysis of nuclear geometry and chromatin organization provides an important complementary approach for investigating the physical requirements of cell-cycle transitions in intact plant tissues.

## Introduction

Cell division is a fundamental biological process that enables cell proliferation, organismal growth, and development while ensuring the transmission of genetic and epigenetic information (Alberts et al., 2002).

From a fundamental physical perspective, one of the major events of the cell cycle is the geometric and structural transformation of chromatin during the G2–M–G1 transition. This process involves extensive chromatin reorganization and the crossing of effective free-energy barriers between interphase and mitotic configurations (Chu & Wang, 2020).

Cell-cycle kinetics may depend on the physical organization and dynamics of chromatin, which vary according to cell identity and function. However, the extent to which the large-scale conformational transformations of chromosomes between interphase and mitosis determine cell-cycle kinetics remains unclear (Contessoto et al., 2026).

In planta, cell division occurs within a developmental context, in cells with established or emerging identities and distinct nuclear and chromatin architectures (Caballero et al., 2024; Chu & Wang, 2022).

Plant nuclei contain euchromatin and heterochromatin, which differ in their structural organization and physical properties. During mitosis, both chromatin compartments undergo extensive, coordinated reorganization to form condensed chromosomes (Pasternak et al., 2023). Their distinct initial configurations may impose different physical constraints on chromosome condensation.

From a molecular perspective, the G2–M transition is regulated by cyclin B–CDK complexes, including interactions involving CDKA and CDKB proteins (Desvoyes et al., 2021; Lee & Torii, 2026).

The dynamic changes in chromatin inside cell cycle in plants were known long time ago (Raynaud et al., 2014), however, these investigations have paid relatively little attention on significant differences in chromatin status/structure in G1 and G2 among cell types.

In many cases, this is may represent an important determinant in cell cycle re-entry (Pasternak/Steimacher, 2026).

Our previous results demonstrated that G2-phase duration in Arabidopsis thaliana is strongly correlated with nucleolar volume (Pasternak et al., 2022). This finding suggests a relationship between nuclear organization and cell-cycle kinetics, raising the question of whether cell-type-specific nuclear architecture contributes to the physical constraints governing the G2–M transition.

Given that different cell types within plant meristems exhibit distinct nuclear architectures and chromatin organization, an important question arises: How do the initial geometric and structural properties of chromatin relate to CYCB1 protein accumulation and the kinetics of the G2–M transition?

CyCB1;1::GFP and CyCB1;1::GUS (Ubeda-Tomas, et al., 2009; Colón-Carmona et al., 1999) lines were widely used as reliable markers of the cell cycle (meristemic) zone in plants. In addition, it is also widely used to determine the quantitative cell cycle activity (level) based on GFP/GUS total intensity per organ (Eloy et al., 2011; Wang et al., 2025). In addition, this marker is also part of a double cell-cycle marker (Hazelwood and Ashraf et al., 2024 for review).

However, our previous observations (Lavrekha et al., 2017) suggest that CYCB1;1 may not quantitatively indicate cell-cycle activity but instead reflect cell-type-specific differences in chromatin organization and timing required for the transition from G2 to metaphase, given the “structural distance” between nuclear size in G2 and in metaphase as described previously (Pasternak and Steinmacher, 2016).

Cell-type-specific nuclear geometry may impose different structural requirements for mitotic chromosome assembly, potentially contributing to differences in CYCB1;1 accumulation and condensin-dependent chromosome remodeling. These observations highlight the need to re-evaluate the interpretation of CYCB1;1 as a quantitative marker of cell-cycle activity.

To address this question, we used a CYCB1;1 reporter to monitor cell-type-specific CYCB1;1 accumulation during mitotic progression and investigate its relationship with nucleus size.

These observations highlight the need to re-evaluate CYCB1;1 not as a quantitative marker of cell-cycle activity, but as an activity linked to nucleus size and the possible speed/duration of the G2-M transition.

## Materials and methods

### Plant material and growth conditions

The plant material used in this study consisted of wild-type *Arabidopsis thaliana* (L.) Heynh. and the transgenic reporter lines CYCB1;1::GUS (Colón-Carmona et al., 1999) and CYCB1;1::GFP (Ubeda-Tomás et al., 2009) all in Col 0 background.

Seeds were surface-sterilized and sown in square Petri dishes containing TK1 medium (Pasternak et al., 2024) supplemented with 1% (w/v) sucrose and 1.1% (w/v) agar (Roth), adjusted to pH 5.6. After sowing, plates were kept at room temperature for 4 h to allow seed imbibition, stratified at 4 °C for 12–14 h, and then placed vertically in a growth chamber at 22 °C under a 16 h light/8 h dark photoperiod, with a photon flux density of 80 μmol m□^2^ s□^1^.

## CYCB1 reporter analysis

### CYCB1;1::GFP analysis

Four-to five-day-old seedlings expressing the CYCB1;1::GFP reporter and grown on solid medium were fixed and counterstained with DAPI. Three-dimensional fluorescence image stacks were acquired as previously described (Pasternak et al., 2015). GFP fluorescence was quantified using a 3D image-analysis plugin to detect and measure reporter signals.

Stained plants were fixed in 4% formaldehyde in microtubule stabilizing buffer (MTSB), washed twice with distilled water for 10 min, incubated in 0.2 mg/l DAPI for 20 min, washed again with distilled water, and mounted on slides with a 120-μm spacer using ProLong Gold Antifade Mountant (ThermoFisher Scientific, Waltham, MA, USA) (Pasternak et al., 2015). In the CYCB1;1::GFP roots, signals were detected after immunolocalization with an anti-GFP antibody following the standard protocol (Pasternak et al., 2015). Confocal images of DAPI-stained samples were recorded using a confocal laser-scanning microscope (LSM 510 META NLO; Carl Zeiss, Oberkochen, Germany) with an LD LCI-Plan-Apochromat 925/0.8 DIC Imm Corr objective. For DAPI excitation, a 740 nm Chameleon laser was used, and emission was detected with a bandpass filter (BP 390–465 IR), while the CYCB1;1::GFP excitation at 488 nm was detected with a bandpass filter (BP 500–550 IR). Serial optical sections were reconstituted into 3D image stacks to a depth of 120 μm, with in-plane (x, y) voxel extents of 0.15 μm and 0.9 μm section spacing (z). Two or three overlapping images (tiles) were recorded for each root. At least 20 roots per variant were scanned.

## CYCB1;1::GUS histochemical analysis

Seedlings expressing the CYCB1;1::GUS reporter were fixed in cold acetone for 20 s and subsequently subjected to histochemical GUS staining. Briefly, seedlings were incubated in sodium phosphate buffer containing 1 mg mL□^1^ X-Gluc at 37 °C for 3–5 h. Following staining, samples were fixed in hot methanol (70 °C), rehydrated in water, and incubated in chloral hydrate solution for 1 h. Samples were subsequently mounted in 50% (v/v) glycerol for microscopic examination.

### Histone marker immunolocalization

Immunolocalization was performed in *A. thaliana* Col-0 seedlings using a previously published whole-mount in situ protocol (Pasternak et al. 2015). The affinity-purified anti-histone H3K4me1 anti-rabbit (Abcam, 1:200 dilution) was used as the primary antibody. The Alexa-488-conjugated anti-mouse or anti-rabbit secondary antibodies were diluted at 1:400.

### Image analysis

Chromatin analysis was performed as previously described (Caballero et al., 2024). Briefly, following 3D imaging, the images were converted to TIFF format, with DAPI, histone-marker, and GFP channels processed separately. Three-dimensional object analysis was performed using the 3D Object Counter plugin in ImageJ (Bolte and Cordelières, 2006). The resulting quantitative data were exported to Microsoft Excel for subsequent analysis. Cell types were identified by converting Cartesian coordinates into cylindrical coordinates and classifying cells according to their radial distance from the root axis.

## Results

### Euchromatin surface area divergence in mitotic zone

In plant nuclei, chromatin can be broadly described as two interacting polymer compartments: euchromatin and heterochromatin. During interphase, euchromatin generally occupies a more spatially extended nuclear compartment, whereas heterochromatin is typically more compact. Consequently, the initial organization of euchromatin may influence the extent of structural remodeling required for mitotic chromosome formation. We hypothesize that differences in euchromatin geometry and nuclear surface area may be associated with differences in the structural requirements of this transition. Quantitative characterization of euchromatin geometry is therefore necessary to evaluate the structural component of this proposed barrier.

To investigate this relationship, we visualized euchromatin-associated chromatin using H3K4me1 immunolabeling (Fig. 1A) and quantified its three-dimensional geometry in root cells with different developmental fates. We found significant differences in euchromatin compactness between outer (epidermal and cortical) and inner (endodermal and pericycle) cell layers (Fig. 1B). To minimize potential effects of differences in DNA ploidy, measurements were restricted to cells undergoing mitosis, identified by the presence of a visible cell plate. Interestingly, despite the cell-type-specific differences in chromatin organization, euchromatin geometry at the mitotic plate was broadly similar across the examined cell files (Fig. 1C).

**Figure 1.**
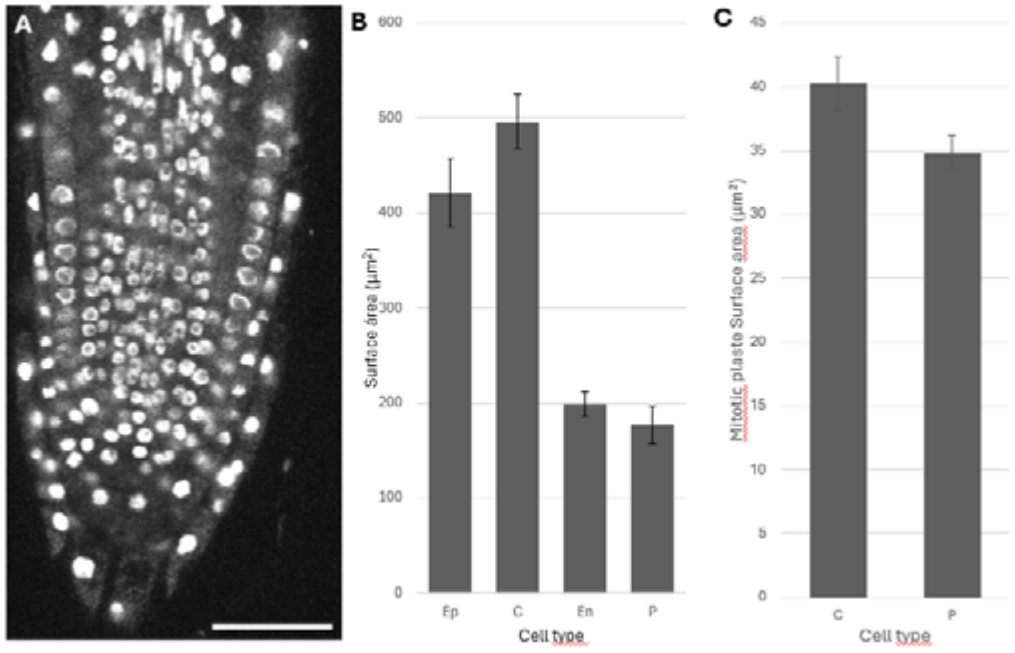
Euchromatin marker geometry in Arabidopsis root. A-H3K4me1 immunolocalization. B - Euchromatin surface area in different cell types; C – Surface area of the mitotic plate. Scale bar – 50 μm.

These observations suggest that distinct cell-type-specific chromatin configurations may converge toward a relatively conserved mitotic architecture. Within our proposed physical framework, the initial structural separation between interphase and mitotic chromatin states may represent an important parameter associated with the remodeling requirements of mitotic entry.

### CyCB1 signal is cell-type specific

CyCB1 is a protein whose activity is detected in late G2 and degraded after mitosis, widely used as a quantitative cell cycle marker. But given the expression window and its dependence on G2 duration, we asked how reliable such data. Figure 2 as well as our previous work (Lavrekha et al., 2017)

**Figure 2.**
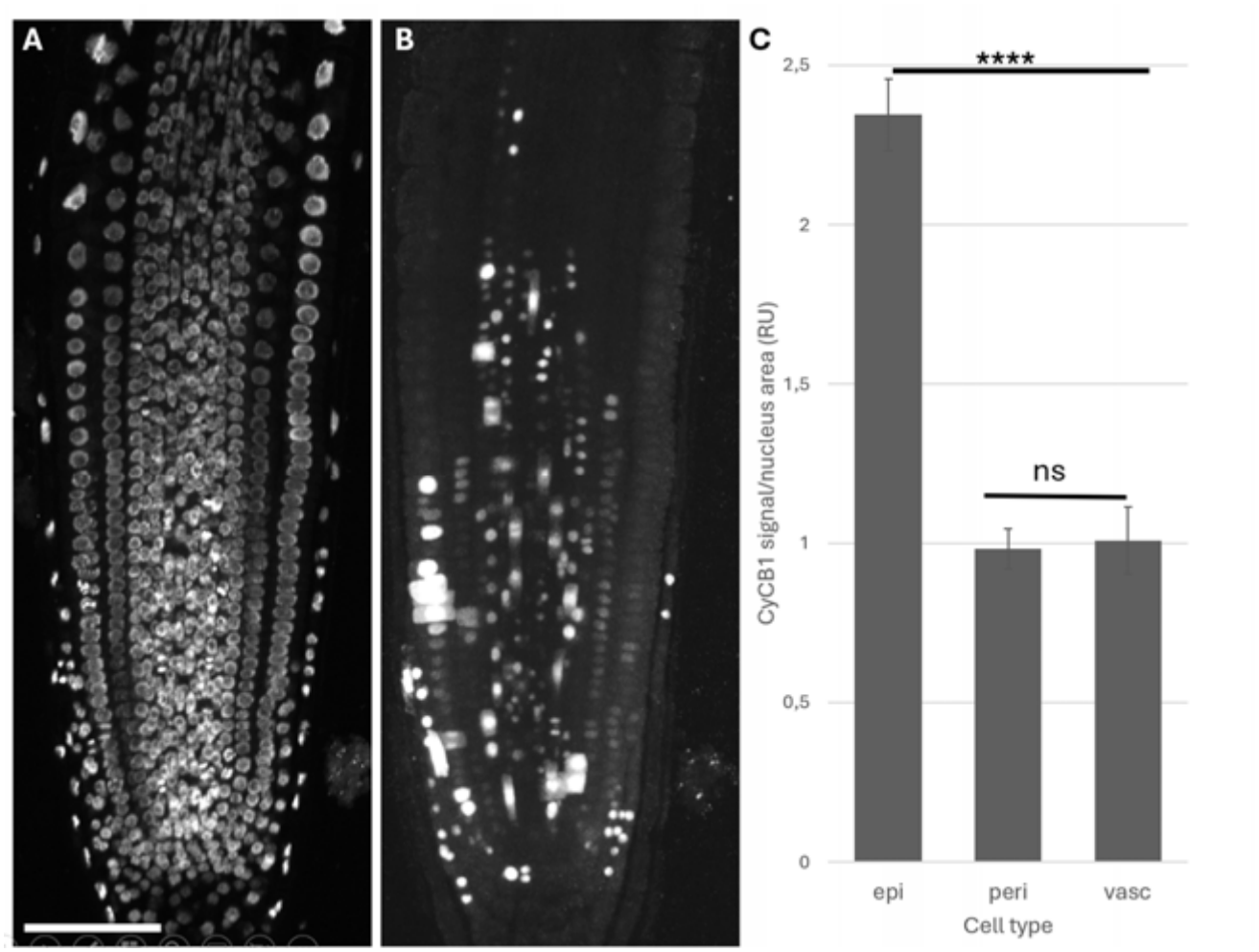
Analyses of cell cycle marker intensity in Arabidopsis root. 5-day-old seedlings were fixed, GFP was detected with an anti-GFP antibody, and co-labeled with DAPI.

**Figure 3.**
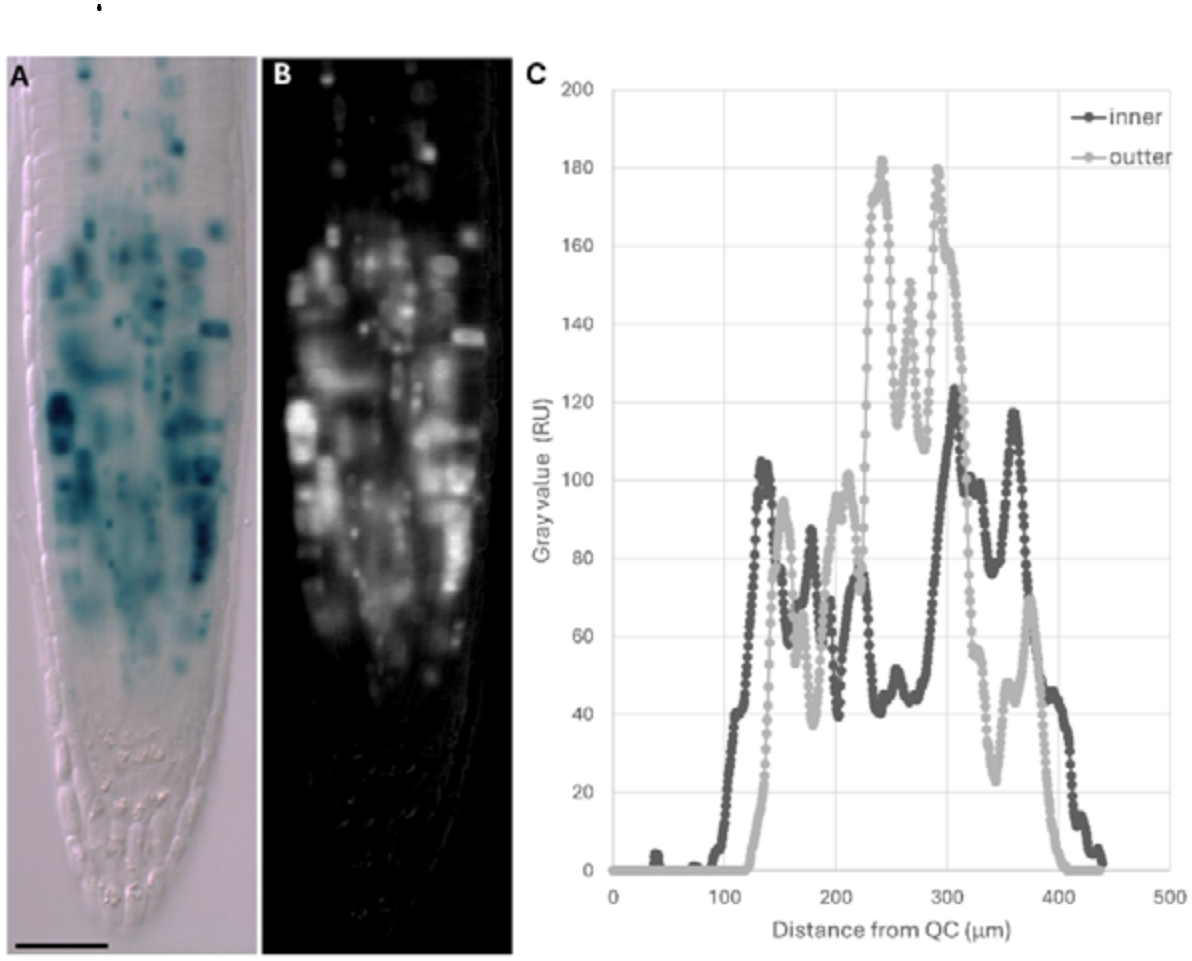
Longitudinal distribution of CYCB1;1::GUS reporter activity in inner and outer tissues of the *Arabidopsis thaliana* root. (A) Representative bright-field image showing histochemical CYCB1;1::GUS staining. (B) Corresponding processed grayscale image used for intensity profiling. (C) Longitudinal intensity profiles obtained from inner and outer root tissues using a 40-pixel-wide line selection in Fiji. Signal intensity is plotted against distance from the quiescent center (QC). The profiles reveal tissue-dependent differences in the magnitude and spatial distribution of CYCB1;1::GUS staining. Scale bar – 50 μm.

A – DAPI; B – CyCB1;1::GFP; C-Quantitative analysis of signal intensity of CyCB1. Epi – Epidermis; Peri – pericycle; Vasc – vasculature. Error bars represent SE. Scale bar – 50 μm.

High CyCB1;1::1 activity is associated with a large nucleus in both GFP and GUS intensity.

The G2-to-mitosis transition in epidermal and cortical cells involves extensive structural remodeling, characterized by a substantial reduction in chromatin-occupied volume, nucleolar disappearance, a marked decrease in segmented chromatin surface area, and increased geometric compactness. Thus, mitotic entry represents a major three-dimensional reorganization of nuclear and chromatin architecture. These changes were particularly pronounced in the more diffuse euchromatin compartment, with significantly greater structural differences between G2 and mitotic states in epidermal and cortical cells than in endodermal and pericycle cells.

Our analysis revealed that these cell-type-specific differences in the magnitude of chromatin remodeling were associated with CYCB1 reporter accumulation and G2 duration. Cells exhibiting greater structural separation between G2 and mitotic chromatin configurations showed higher CYCB1 reporter signals and longer G2 duration. Conversely, cells with smaller structural differences exhibited lower CYCB1 reporter signals and shorter G2 duration. These findings establish an association between chromatin geometry, CYCB1 reporter accumulation, and cell-cycle kinetics across different root cell types.

Importantly, extensive chromatin remodeling does not necessarily prevent mitotic progression, as the mitotic machinery actively mediates chromosome condensation and nuclear reorganization. Therefore, the geometric distance between G2 and mitotic chromatin configurations should be interpreted as a measure of the structural transformation associated with mitotic entry, rather than as an absolute barrier to cell-cycle progression. We propose that the initial chromatin architecture represents an upstream physical condition that may influence the remodeling requirements and kinetics of the G2-to-M transition.

## Discussion

Late G2 is generally characterized by an enlarged nucleus, whereas mitotic entry in-volves chromatin condensation into distinct mitotic chromosomes, accompanied by extensive reorganization of the nuclear envelope and nucleolus. Collectively, these coor-dinated changes can be described as a structural nuclear transition. Within a given plant species, cells with the same chromosome complement may exhibit relatively similar total mitotic chromosome volumes despite substantial differences in their interphase nuclear architecture.

However, the physical requirements of the G2–M transition cannot be characterized by chromatin volume alone. Nuclear geometry, surface area, chromatin topology, and polymer organization may impose additional constraints on chromosome condensation. In particular, the transformation of an enlarged G2 nucleus into compact, individualized mitotic chromosomes requires extensive three-dimensional chromatin remodeling, co-ordinated with nuclear envelope disassembly and nucleolar reorganization. Consequently, cell-type-specific differences in initial nuclear architecture may influence the structural requirements and kinetics of the G2–M transition (Ma et al., 2015).

Additional evidence linking CYCB1;1 accumulation to delayed cell-cycle progression comes from studies of DNA-damage responses, in which increased CYCB1;1 expression or reporter accumulation has been observed under conditions associated with cell-cycle arrest or delay (Schnittger and De Veylder, 2018). These observations indicate that elevated CYCB1;1 reporter levels do not necessarily correspond to increased cell-division frequency.

Taken together, our quantitative analysis revealed a correlation between nuclear geometry, CYCB1;1 reporter accumulation, and cell-type-specific G2 duration. Cells with larger nuclei and greater structural remodeling requirements exhibited higher CYCB1;1 reporter accumulation and longer G2 duration. These findings support a close relationship between chromatin geometry and cell-cycle kinetics and suggest that CYCB1;1 reporter accumulation should be interpreted in the context of nuclear architecture rather than as a direct measure of cell-division activity alone.

## Conclusions

Our findings support a model in which cell-type-specific nuclear architecture influences the structural requirements of the G2–M transition. Greater differences between interphase and mitotic chromatin configurations may require more extensive remodeling, potentially contributing to differences in CYCB1;1 accumulation and cell-cycle timing. Direct measurements of remodeling dynamics and CDK activity are needed to establish the underlying mechanism.

## Declaration of AI usage

During the preparation of this manuscript, the authors used ChatGPT 5.0 in order to correct language errors. After using this service, the authors reviewed and edited the content as needed and take all responsibility for the content of the published article.

